# Drop-by-drop Addition of Reagents to a Double Emulsion

**DOI:** 10.1101/2024.05.16.594444

**Authors:** Thomas W. Cowell, Wenyang Jing, Hee-Sun Han

## Abstract

Developments in droplet microfluidic assays have facilitated an era of high-throughput, sensitive single-cell, or single-molecule measurements capable of tackling the heterogeneity present in biological systems. Relying on single emulsion (SE) compartments, droplet assays achieve absolute quantification of nucleic acids, massively parallel single-cell profiling, identification of rare variants, and more. Double emulsions (DEs) have seen new interest in recent years for their potential to enable new droplet assays and build upon SE techniques. DEs are compatible with flow cytometry enabling high-throughput multi-parameter drop screening and eliminate content mixing due to coalescence during lengthy workflows, addressing inherent limitations of SEs. Despite these strengths, DEs lack important technical functions that exist in SEs such as picoinjection or any other method for adding reagents to droplets on demand. Consequently, DEs cannot be used for multistep workflows which has limited their adoption in assay development. Here, we report a simple device achieving picoinjection of DEs. We developed strategies to enable active manipulations on DEs by converting DE inputs to SEs on chip. The released aqueous cores of the DE can be manipulated using existing SE techniques, such as reagent addition, before reforming a DE at the outlet. We identified device designs and operation conditions achieving drop-by-drop reagent addition to DEs and used it as part of a muti-step aptamer screening assay performed entirely in DE drops. This work enables the further development of multistep DE droplet assays.

## Introduction

The field of droplet microfluidics uses monodisperse emulsions as individual micro-scale reaction vessels, enabling a diverse suite of applications spanning highly-parallelized biological assay, sensitive single-molecule detection and quantification, or the synthesis of functional microparticles.^1–10^ Due to their simplicity, water-in-oil single emulsions (SEs) are the most widely used droplet format.^11^ Although microfluidic production of higher-order emulsions was established concurrently, both device fabrication and operation was more difficult for multiple emulsions which limited their wider use.^12–14^ In applications like drug delivery, where multi-layer emulsions mold unique 3-d structures onto the micro-particles or capsules they produce, the increased complexity of higher-order emulsions was justified.^15–17^ But when droplets are used merely as compartments or reaction vessels, increased levels of emulsification was seen as excessive or unnecessary.

This view has shifted over time as unique advantages of water-in-oil-in-water double emulsions (DEs) have emerged and improvements in DE devices have reduced the complexity of drop production.^18–20^ DEs achieve highly parallelized compartmentalization akin to SEs while being suspended in an aqueous medium. Consequently, DEs have been shown to be compatible with broadly accessible flow cytometry instruments.^21–26^ When compared with equivalent SE droplet screening platforms, DE-flow cytometry offers a substantial improvement in terms of availability, functionality, and ease of use. SE droplet screening requires a custom-built sorting stand and significant microfluidic experience to achieve 1-3 color fluorescence detection and ~200 Hz sorting speeds.^27–29^ In contrast, flow cytometry instruments, which are commonly featured in shared facilities and easy to use, achieve >10 color fluorescence detection, and higher droplet sorting speeds.^30,31^

The unique structure of DEs provides other advantages beyond flow screening. Material transport across the oil shell has been applied for small molecule diffusion following an in-drop reaction to interface incompatible steps or for nutrient exchange to enable long-term cultivation of microbes.^32,33^ Further, DEs are robust against coalescence.^20^ In response to stresses, SE compartments merge with each other and combine, which results in reduced monodispersity and a loss of single entity resolution. To limit the negative impacts of coalescence, most commonly used SE-based bioassays limit in drop operations to a single incubation or thermocycling step prior to pooling or analysis.^34–38^ DEs can address this limitation, as they release drop contents into the outer phase during coalescence. By releasing into the outer phase, any further reactions are halted, mixing between drops is eliminated, and the remaining DE keeps its monodispersity.

These properties make DEs ideal for multistep workflows, which rely on monodisperse drops throughout all stages of a workflow. However, DEs lack methods to add reagents on demand and so they have not been able to realize this potential for robust multistep workflows. In SEs, fresh reagents can be added to drops using picoinjection^39^ or drop merging,^40^ which add a controlled volume of reagents to each drop. However, applying these approaches directly to DEs is challenging as the additional emulsion layer prevents direct access to the inner aqueous compartment. In a SE picoinjector, the simplest method of reagent addition, each drop comes in contact with a stream of reagents that are to be added. The two fluids are separated by a single surfactant-stabilized interface which is disrupted by nearby electrodes causing them to fuse.^41^ Ultimately, picoinjection forms a new drop consisting of the original drop contents and a controlled volume of added reagents.^40^ Due to the differences in drop structure, directly implementing picoinjection or other reagent addition methods has not been possible in double emulsions.

Instead, progress towards sequential or time-delayed reactions in DEs has focused on pre-loading the DE with multiple internal compartments during production.^42–46^ In response to an electrostatic,^42,43^ thermal,^44^ optical,^45^ or chemical stimulus^46^ the contents of the segregated compartments are mixed and combined to trigger a subsequent reaction in the droplet. This approach is severely limited for two reasons. First, multi-core DE production is more challenging both in terms of device fabrication and operation. Glass capillaries are used most often but struggle to produce drops that are small enough for use with FACS, which requires sizes <100 µm.^47,48^ PDMS devices can make smaller drops but require more complex multi-layer fabrication and precise flow synchronization to produce multi-core DEs. Second, preloading of components is not suitable for most multistep molecular biology reactions as it requires all components to be subjected to a single shared temperature profile. Many enzymes and reagents are sensitive to thermal degradation or lose activity over time. Addition of fresh reagents is a key requirement for robust multistep workflows, which has not yet been realized in DEs.

Here, we report a new microfluidic approach that achieves on demand reagent addition to DEs expanding workflows to include multistep reactions. The newly designed device converts DE inputs into SEs on chip, so that each drop is accessible for picoinjection. After adding fresh reagents, the same device reforms the DE achieving on demand, high throughput reagent addition. We then demonstrate a two-step DE workflow to generate a library of aptamer drops then screens and recovers drops that display light-up fluorescence. This assay leverages the unique advantages of DEs across two steps enabled by reagent addition. In the first step, each DNA template in the aptamer library is digitally amplified in drops, while the DE format ensures robust compartmentalization avoiding thermal-induced merging. Next, RNA synthesis reagents are added, avoiding thermal degradation of these temperature sensitive reagents. The reformed DE again maintains compartmentalization during the lengthy RNA synthesis step and ensures uniform aptamer yields across drops. Further, the DE output is compatible with flow cytometry enabling high throughput screening. If this assay were performed in SEs, drop merging could confound the aptamer screen as larger merged drops could produce higher signals that are not correlated to the encoded sequences due to an increased aptamer yield or dye amount. As shown, these new strategies enable multistep assays in DE droplets, achieving a substantial improvement in compartmentalization and droplet screening when compared with existing SE assays.

## Results and Discussion

### Design Principles for active manipulation of DE droplets

Our approach to reagent addition in DEs consists of a series of modular operations implemented in a single microfluidic device (Scheme 1). The first operation introduces a close-packed DE through reinjection and respaces the drops so that each travels one-by-one. The next operation converts each respaced drop into a SE through controlled release of the aqueous core. Following DE to SE conversion, the next operation adds a controlled volume of reagents to each droplet by picoinjection. The final operation forms a high-quality DE output from these droplets. The microfluidic device implementing all these functions is shown in Figure S1A. Performing picoinjection requires a fluorophilic surface treatment, while DE formation necessitates a transition to a hydrophilic surface treatment. Co-patterning of fluorophilic and hydrophilic surfaces was achieved using a modified sequential flow-confinement strategy consisting of polyelectrolyte deposition and Aquapel treatment (Figure S1B-C). This surface treatment strategy enables the single-layer device to implement all the constituent operations that achieve reagent addition to a DE. The following sections detail the design and operational constraints for each of the modular operations.

### Reinjection of Double Emulsions

After performing the first assay step in DE droplets, the emulsion is reinjected into a second device for drop-by-drop manipulation. During reinjection, drops need to be delivered to the device “closed packed” so that they can be respaced to travel one-by-one. Due to the differences in emulsion properties, the existing methods of SE reinjection^34^ must be modified for DEs. First, the aqueous phase is less dense than the fluorinated phase, which causes SEs to float in their continuous phase while DEs sink. Accordingly, the syringe needs to be inverted for the DE to pack under gravity. A detailed discussion of preparing and delivering a close packed DE is included in the supplementary text. During SE reinjection, the inlet surface is treated with a fluorophilic coating to maintain the drop stability during injection. For DEs, the equivalent hydrophilic surface treatment facilitates stable reinjection. However, this requires 3 separate surface treated regions (hydrophilic-fluorophilic-hydrophilic) which is challenging to implement using flow confinement and would increase the complexity of device fabrication.

As an alternative, we found that when the device height is sufficiently greater than the DE core diameter, the emulsion stability was not strongly influenced by the surface wetting properties of the inlet. A systematic characterization of DE stability during reinjection into a fluorophilic inlet is shown in Figure 1A. We express the size of the various DE inputs as the ratio of the DE core diameter to the inlet height. Interestingly, we found that the oil shell diameter does not impact reinjection behavior. For core diameter sizes less than 70% of the inlet height, the DE remained stable during reinjection (Figure 1B). Above this critical size, the DE is squeezed against the channel surfaces causing uncontrolled coalescence of the various phases (Figure 1C). To facilitate reinjection of larger sized DEs, an additional hydrophilic surface treatment could be applied to the inlet, or a multi-layer device could be used so that the inlet height is greater than the remainder of the device. Our single-layer design keeps the device complexity low while accommodating a large range of DE inputs.

**Figure 1:**
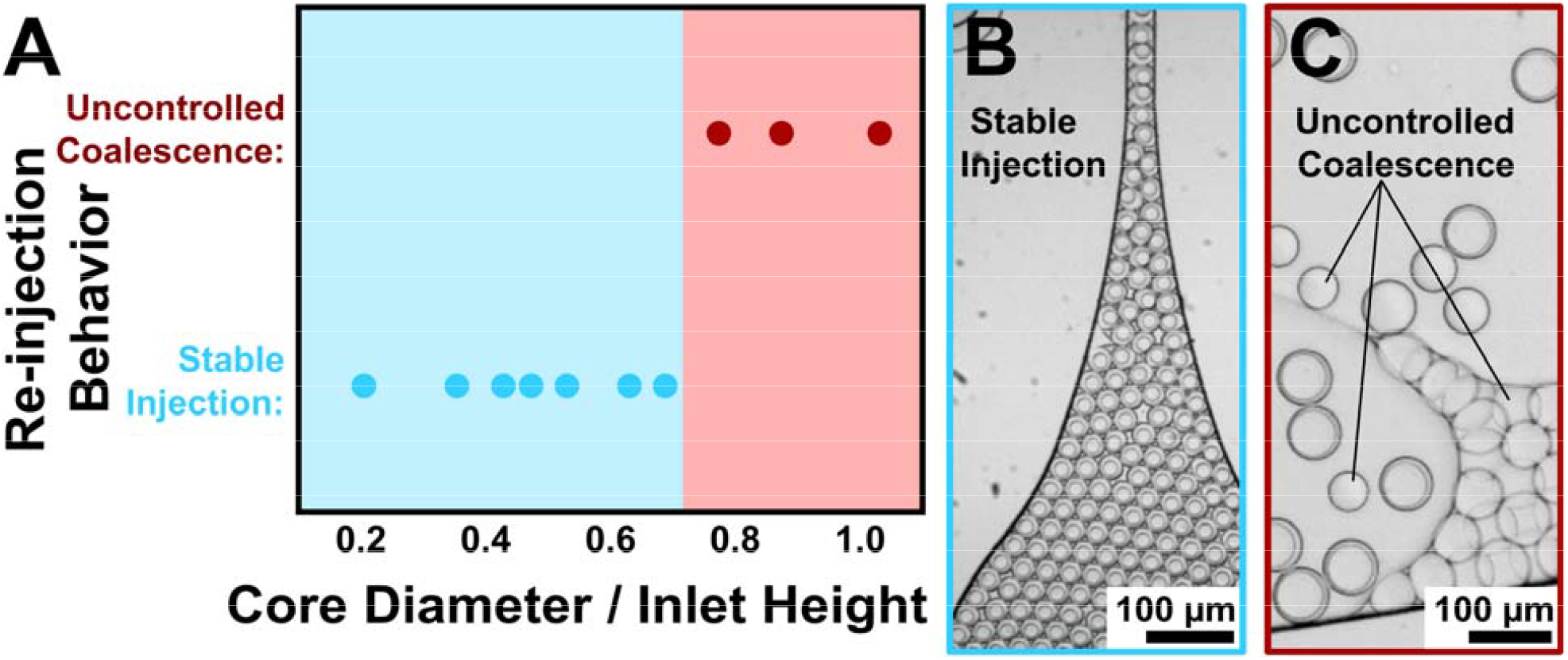
A) Characterization of suitable DE sizes for successful reinjection relative to the device height at the inlet. B) Microscope image showing stable reinjection of a DE. C) Microscope image showing uncontrolled coalescence during reinjection of a DE to a fluorophilic inlet.

Following reinjection, the inlet geometry funnels the drops into a single file for respacing. For single emulsion methods, respacing is achieved using additional oil that separates each drop so that they travel one-by-one. In SE devices, SE respacing produces a regular frequency of spatially separated SE drops. Although DEs can be respaced in their continuous phase in the same way, the picoinjection channel requires the oil-phase as the continuous phase. Consequently, we must “respace” the close packed DE in fluorinated oil which further emulsifies each drop to form a water-in-oil-in-water-in-oil triple emulsion (TE). We refer to these TEs as respaced DEs.

### Conversion of Respaced DEs to SEs by Controlled Core Release

Following reinjection and respacing, the next step makes the inner aqueous compartment of each drop accessible for picoinjection. This was achieved using a constricted channel that reliably releases cores by rupturing each respaced DE. To characterize the range of suitable sizes for core release in this device, various DE inputs were screened (Figure 2A). The input drop sizes are expressed as a ratio of the DE core diameter to the constricted channel width. As described in the previous section, the inlet height during reinjection constrains the maximum size of DE inputs for this device to ~2 times the constriction width. DEs above this size would likely release in the constriction but could not be tested as they coalesce in the inlet during reinjection. Between this upper limit and ~1.2 times the constricted channel, DEs reliably converted to SEs. Figure 2B shows a series of sequential images during controlled core-release. In the constricted channel, the oil shell is sheared until the TE ruptures. This releases the inner core and the outer aqueous (OA) pocket as single emulsion drops. This controlled core release process results in each DE in the input converting to a SE one-by-one making them accessible for picoinjection.

**Figure 2:**
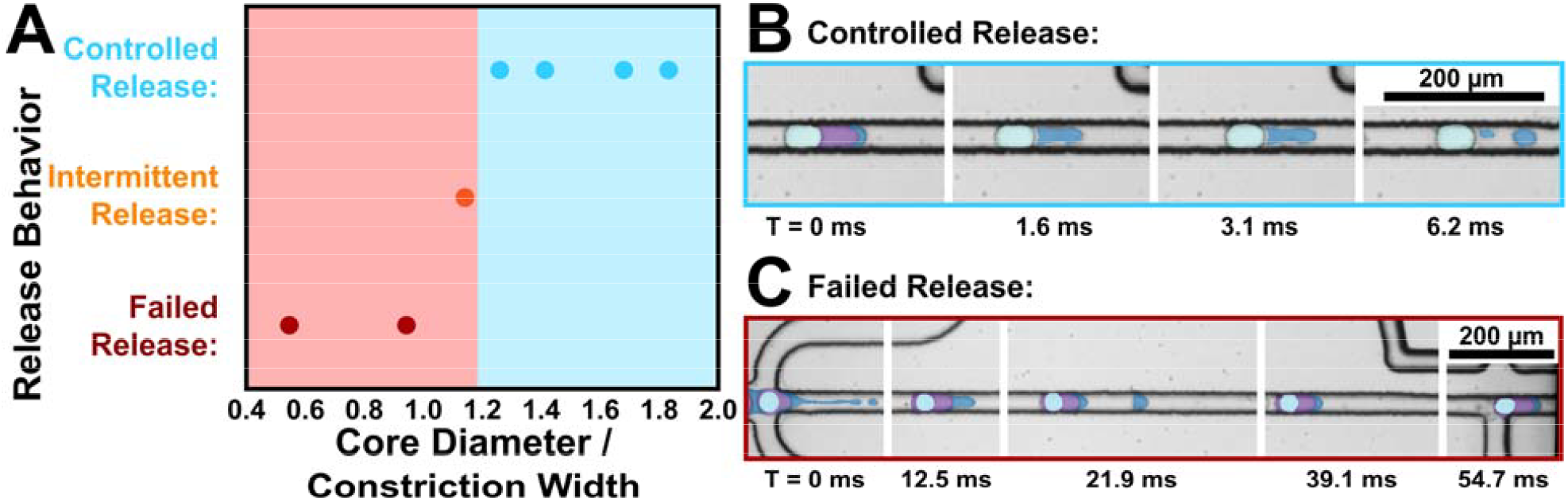
A) Characterization of suitable DE input sizes for core release behavior. B) Sequential images showing controlled release of a DE as it releases its core as a SE. C) Sequential images showing failed release as a respaced DE passes through the constricted channel without conversion to SE. Input cores are light blue. The oil shell is labeled purple. The outer aqueous phase is dark blue.

As the size of the DE input is decreased further (~1.1), the emulsion switching process becomes unreliable and not all DE cores are released (orange dot in Figure 2A). Below this size, the emulsion would fail to release any cores as it passes through the constricted channel (Figure 2C). Previous work has indicated that active electrodes can facilitate the rupture of TEs.^49^ However in this device configuration, including active picoinjection electrodes could not induce TE to SE conversion in cases where passive release failed. This may be explained by the substantially greater oil shell thickness in our system. Regardless, a large range of suitable DE inputs were identified that can be controllably converted drop-by-drop to an accessible SE format. The design of the inlet and constricted channel could be rescaled to accommodate DEs of other size ranges.

### Picoinjection of Released DE Cores

The released cores from each DE can be picoinjected using standard SE manipulation methods.^39^ At this stage, the picoinjected drops could be collected as a SE to achieve a two-step workflow where the first step is performed in a DE and the second steps is performed as a SE. The picoinjector could also be replaced with any other SE manipulation module to implement other SE manipulations downstream of an initial DE droplet step. However, to fully leverage the advantageous properties of DEs, we want to collect the drops as a DE after manipulation.

DE production from SEs is not trivial, especially when the SE stream is not regularly sized or spaced as is the case here. Since we want to ensure that every released core is picoinjected, the reagent flow rate is high relative to the droplet frequency. Consequently, the device alternates between picoinjection of cores and production of reagent-only drops. Since these drop types are not identically sized, they travel at different speeds such some drops are touching by the time they reach the DE reforming junction. Accordingly, the resulting stream of SEs has an irregular frequency. In traditional DE production devices, which adopts a sequential emulsification method, relies on synchronization of drop arrivals with the DE pinch-off to form a monodisperse DE. Since this approach is not applicable to irregular droplet stream as in our case, further investigation into DE production methods are needed to recover a DE output.

### Reforming a High-quality DE Output after Picoinjection

Our previous work described a type of DE drop generation in a confined, one-layer device that was largely self-synchronizing due to core-filling induced DE pinch-off.^20^ However, this study used an upstream drop maker which produces a regularly sized and spaced stream of SE drops. Here we extend these core-filling strategies to generate DEs from irregular streams of SEs. For functional bioassays, the DE formation step must meet the following requirements. Each picoinjected core should be loaded by itself into a single DE drop. It is important that cores are not split across multiple DEs or loaded along with other drops as a multi-core DE. The presence of empty oil drops is not important to DE quality as these oil drops can be ignored in most cases or can be separated by centrifugation when necessary. With these criteria, we can assess device performance by evaluating DE outputs across a range of operational parameters.

Although there are many parameters that influence DE production, including flow rates, fluid properties, surface treatments, and device dimensions, most of them are not freely tunable. Each of the upstream flow rates are used to control respacing and picoinjection. The viscosity, surface tension, and composition of each fluid component is set by the application. The surface treatments were selected for SE manipulation and DE production. The device height was fixed to enable DE reinjection. The junction width was chosen to make FACS-compatible DEs with total sizes below 100 µm. These restrictions leave the OA flow rate as the remaining handle for tuning DE production.

To test if the OA flow rate is sufficient to control DE production, we used the device to form DEs using SE inputs that span the relevant range of drop sizes identified in previous sections. To achieve this, monodisperse SEs were produced using a separate device and reinjected. The upstream flow rates were held constant, while the OA flow rate was varied, and the quality of the resulting DE outputs was characterized (Figure 3A). Drop sizes are expressed as a ratio of their circular cross-section to that of the rectangular junction cross-section. From this characterization of DE production, we identified a large range of drop sizes (0.30 to 0.87) where there was some range of flow rate capable of generating a high-quality DE (Figure 3B). Outside this broad range of suitable sizes, some issue with DE production could not be addressed using the OA flow rate. At small SE sizes (<0.30), formation of multi-core DEs unavoidable (Figure 3C). Increasing the OA flow rate reduces the average number of cores per drop, but the inner stream began to jet before a fully single-core loading could be achieved. At large SE sizes (>0.87), aqueous cores do not easily fit within a single DE drop, resulting in core-splitting even at low flow rates (Figure 3D). Further reductions in the OA flow rate resulted in oil-phase jetting that was unsuitable for DE production.

**Figure 3:**
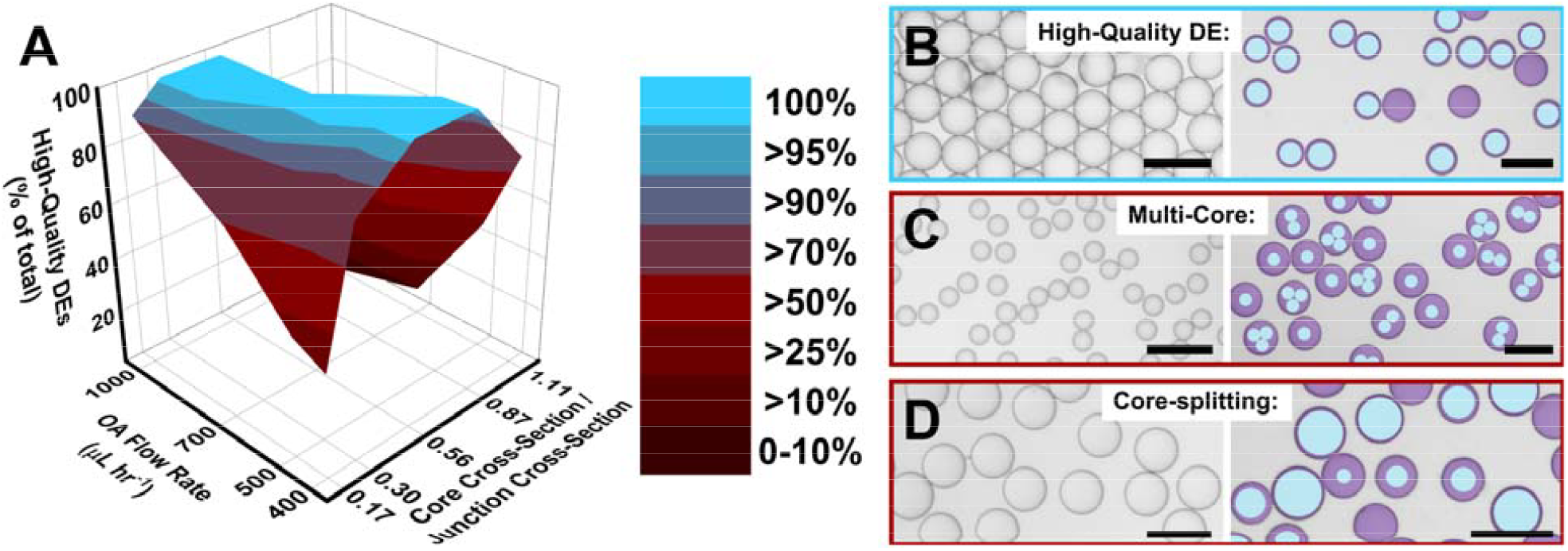
A) Characterization of the DE formation across a range of operational parameters. Height and color indicate the fraction of drops produced that are a high-quality DE. B-D) Images showing a SE input (left) and the resulting DE output (right) to illustrate the key operational modes for DE formation. Scale bars are 100 µm. Within a broad range of sizes, the flow can be optimized to achieve high-quality DE production. At small sizes, multi-core loading results reducing DE quality. At large sizes, drops do not fit within a single DE. Core-splitting can occur causing a low-quality output. DE cores are labeled light blue, and oil shells and excess oil drops are purple.

Importantly, within the identified operational regime, each OA flow rate can accommodate a smaller but still broad range of SE sizes. This is the case because the DE reforming junction operates in a flow regime that is largely self-synchronizing. Each aqueous core contributes a substantial fraction of the volume required to initiate pinch-off of the filling oil droplet. As such the arrival of each aqueous core effectively triggers the DE droplet to detach. This phenomenon facilitates single core DE formation on the irregular drop frequencies, polydisperse inputs, and even in cases where SE drops are in contact with adjacent drops (Supplementary Video 1). As we show here, it is possible to reliably reform high-quality DEs from challenging SE streams.

### Reagent addition to a Double Emulsion

We evaluated the combined performance of the device and assessed the quality of the resulting DE after reagent addition. If the process is successful, every aqueous core from the input DE will have a controlled volume of reagents added resulting in a larger, monodisperse, single core DE output. The presence of excess oil drops or reagent-only drops do not impact DE-based bioassays and should be excluded from assessment of DE quality. To clearly distinguish the composition of aqueous drops in the DE output, we fluorescently labeled the inner aqueous phase of the input DE and the reagent stream with distinct dyes. The oil and outer aqueous were not fluorescently labeled but can be readily distinguished. Figure 4A-B shows the device during operation. The picoinjector alternates between injection of released cores (green drops) and production of reagent-only drops (yellow drops). Because these drops are different sizes they catch up and travel as a pair that is separated at the junction to form single core DEs (Supplementary Video 2-3). Figure 4C-D shows the DE before and after addition of reagents. The DE output also contains excess oil drops and reagent-only droplets. Since reagent only drops will not be positive in any assay, they do not impact performance. Likewise, oil drops can be ignored in most applications or separated by density. The components of the emulsions in Figure 4 are colored to improve the visual clarity by using fast-camera recordings (A-B) or fluorescent signals (C-D). The histogram of drop sizes (Figure 4E) shows that output DE is single core and monodisperse (6.65%), achieving uniform addition of reagents to each drop in the input. In this configuration, the added reagents constitute approximately two thirds of the final drop volume which is typical for a picoinjector of this type and is sufficient to implement most multistep reactions. Modifying the flowrate of the reagent stream adjusts the volume added to each drop.^50^ These results confirm our device can achieve all the key requirements for reagent addition in DEs.

**Figure 4:**
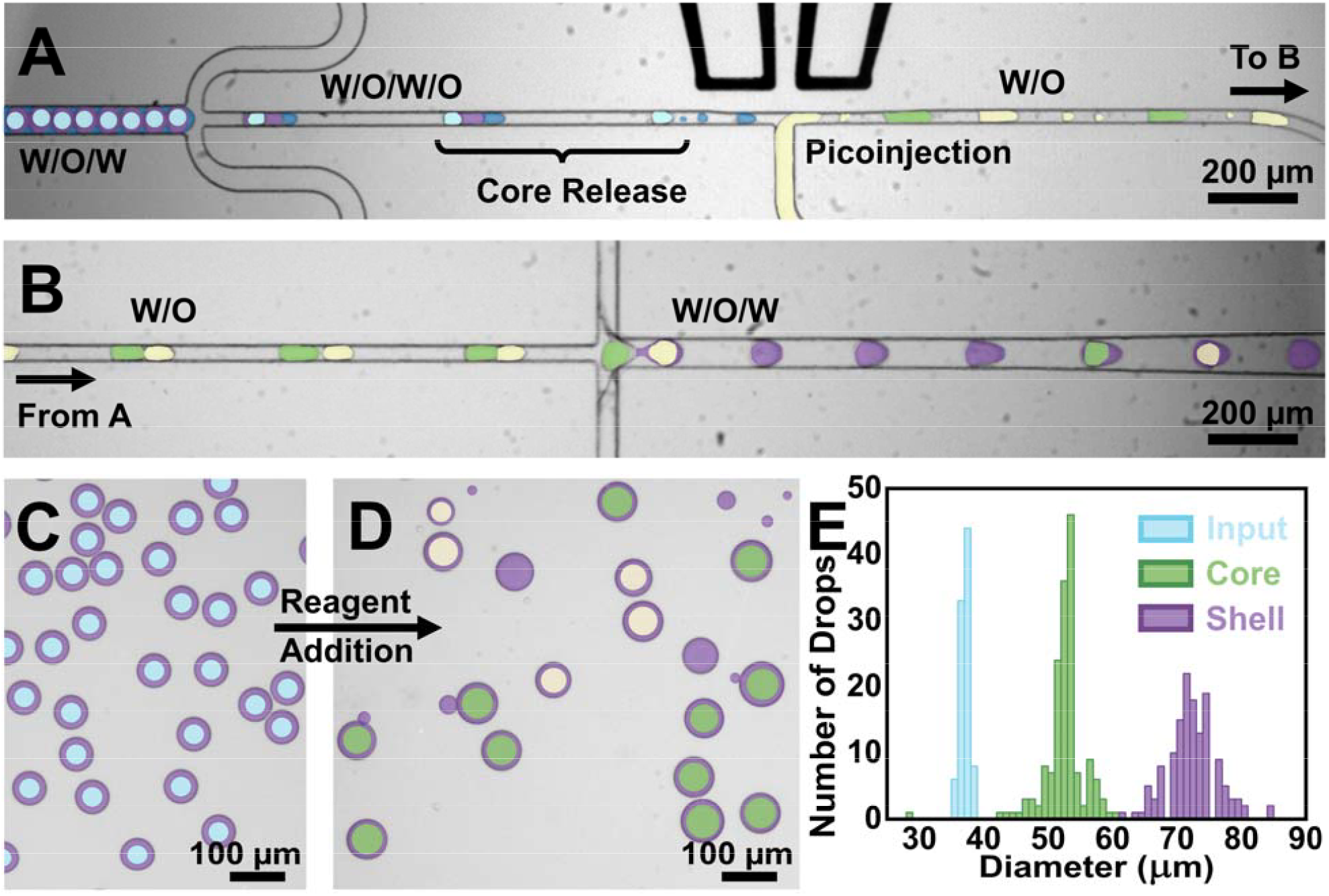
Operation of the DE picoinjection device during drop-by-drop reagent addition to a double emulsion. Input cores are light blue. Excess oil drops and oil shells are labeled purple. The outer aqueous phase of the input is dark blue. The reagent stream and reagent-only drops are yellow. Picoinjected cores are labeled green. A) Microscope image showing respacing, core-release, and picoinjection. B) Downstream image showing DE production. C) Image of the input DE. D) Image of the output DE. E) Histogram of drop sizes.

### Implementing a multistep aptamer library screen in Double Emulsions

With this newly developed device, we performed droplet screening of light up RNA aptamers which requires two distinct steps entirely in DEs. RNA light-up aptamers have attracted interest for their ability to label intracellular RNAs enabling localization and tracking in live cells.^51–53^ Upon aptamer binding to the dye, it “lights up” becoming fluorescent. DFHBI dyes are of particular interest with their capability to facilitate live-cell imaging due to low cyto-toxicity, high membrane permeability, and fluorescence properties that closely match GFP.^54^ There is considerable interest in identifying new light up aptamers for multiplexed labeling. However, light up aptamer screening can be challenging requiring each sequence variant to be compartmentalized and assessed for their capability to turn on the fluorescence of dyes at extremely high throughput.

A standard aptamer screen can be broken into the following steps: expression of a diverse aptamer library, functional assessment of each variant, and enrichment of high functioning sequences. Particle display methods enable assessment and sorting of aptamer-induced fluorescence using flow cytometry.^55^ Using DEs instead of microparticles offers a unique ability to screen aptamer properties that are not directly associated with their binding ability. Since drop assays confine reaction products within the compartment,^56^ binding strength is decoupled from the screened property, in this case light-up fluorescence.

DE properties are useful throughout all steps of a droplet aptamer screen. In the first step, DEs solve PCR merging issues ensuring a monodisperse emulsion that effectively compartmentalizes each unique sequence. Maintaining the monodispersity of droplets is also critical for the pico-injection step as it maintains the uniform spacing between drops upon reinjection and respacing, which is critical for drop-by-drop manipulation. Since we use a DE also for the second step, the uniform size of drops is perfectly maintained during incubation resulting in the uniform aptamer yields across drops. Lastly, the use of DEs enables high-throughput droplet screening by flow cytometry which is an efficient method of screening the aptamer library. Importantly, this workflow is generalizable and could be applied to screen light-up aptamers resulting in improved fluorescence properties or distinct emission spectra or to target other aptamer functions.^57^

The DE aptamer screening workflow implemented here is shown in Scheme 2. Using the newly developed reagent addition device, we introduced fresh reagents to each DE after the PCR step completes enabling a multistep workflow entirely in a double emulsion. For our screening experiment, we generated the library yielding ~10^6^ unique sequences by modifying the broccoli sequence to contain 10 consecutive random bases within the dye-targeting region (Table S1). In the first step, this library of unique DNA oligos are loaded into drops for digital PCR amplification. The loading efficiency was set at 10-15% ensuring most drops contain 0 or 1 template copies. In this experiment, the known broccoli sequence was added to the pool at a rate of ~1 in 5,000 to reduce sorting times. The post-PCR DE remains highly monodisperse (Figure 5A) confirming that the DE maintains single template compartmentalization and ensures uniform yields across drops.

**Figure 5:**
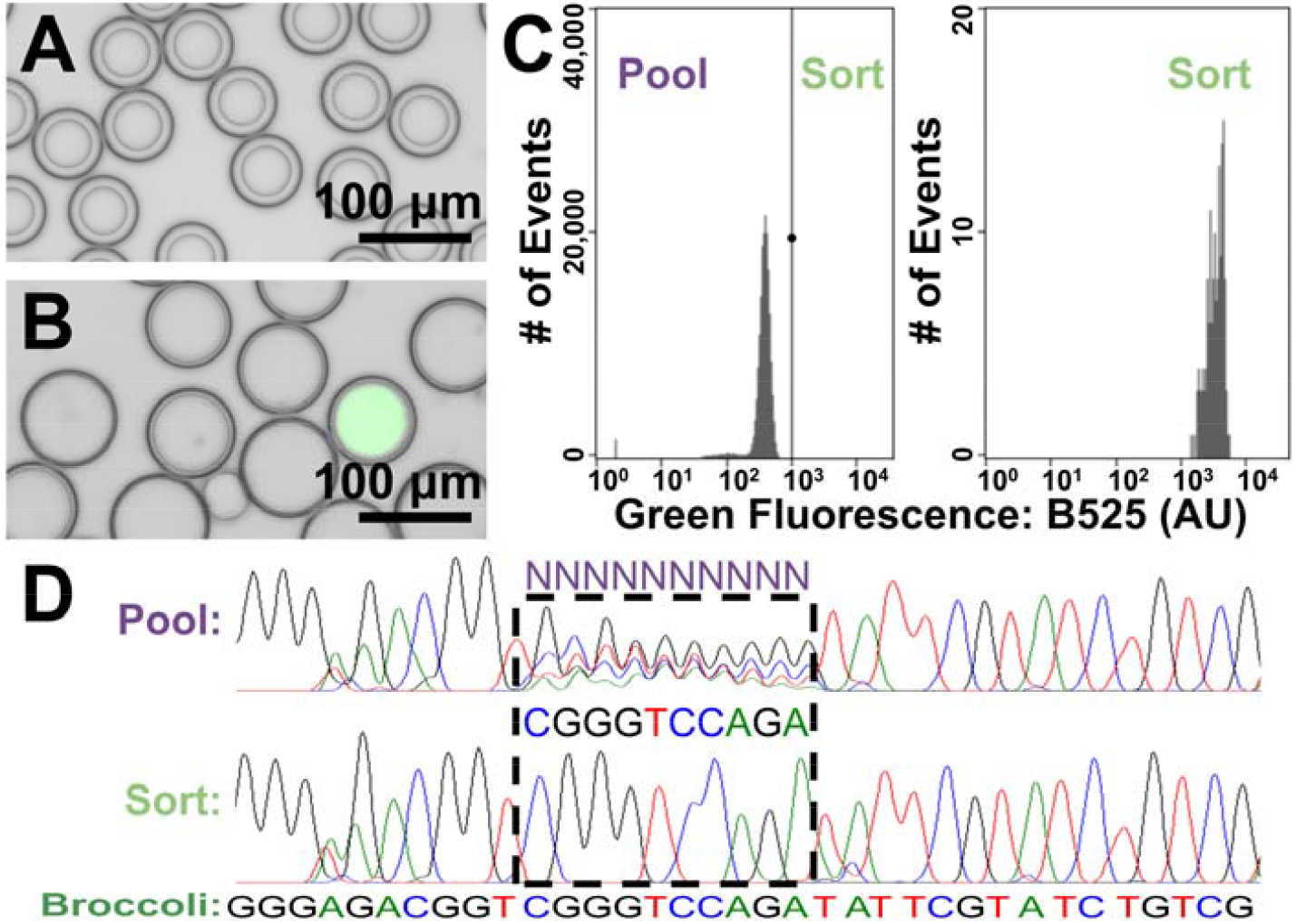
A) Microscope image of the DE after completion of the PCR step. B) Composite image (brightfield and GFP) showing the DE after RNA synthesis and staining with the target dye. Some drops with elevated fluorescence were observed. C) Fluorescence histogram resulting from flow screening of the DE aptamer library. Drops displaying elevated fluorescence above the threshold were sorted. D) Sanger sequencing results from the aptamer screen. Sorting recovers the correct broccoli sequence.

Next, the DE was reinjected and processed to add reagents for RNA synthesis generating a new DE output for the second step (Figure S7A). These drops were incubated overnight to produce aptamers. DFHBI dyes were added to the OA phase staining each droplet by diffusion across the shell. This ensures that the workflow can proceed even in cases where target dye is inhibitory. After staining, a subset of the emulsion displayed significantly increased green fluorescence due to successful PCR amplification and T7 RNA synthesis of functional aptamer sequences (Figure 5B). Using a low-pressure, microfluidic flow-sorter (WOLF G2) which can process larger sized DEs, the pool of droplets was screened and sorted to recover the drops that were fluorescence positive (Figure 5C). DNA amplicons from these drops were recovered and sequenced (Figure 5D). The unsorted pool lacks a consensus sequence resulting in consecutive Ns, while the DE sort successfully recovers the correct Broccoli aptamer sequence. We further assessed the fluorescence properties of the pre- and post-sort aptamers. Equal aptamer concentrations of each condition were prepared and assessed for their light up fluorescence in the presence of the target dye. As expected, a significant increase in emission at 501 nm was observed for the sorted aptamers relative to the unsorted pool (Figure S7B). These results confirm the functionality of the multistep DE screening platform.

## Conclusion and Perspectives

Here, we report a microfluidic approach achieving on-demand reagent addition in DEs. Our modular device functions by reliably releasing each of the individual aqueous cores from an input DE so that they are accessible to conventional drop-by-drop SE manipulation techniques. Although we focus on implementing and optimizing a picoinjector in this work, the principles of DE to SE conversion can be applied to facilitate any other SE manipulation technique by replacing the picoinjection module with a pairwise drop merger,^58^ drop splitter,^59,60^ drop sorter,^34^ etc. In this way, SE manipulations can be applied generally on DE inputs.

Importantly, we also developed DE reforming strategies enabling on demand emulsion switching by converting SEs back to DEs. We characterized device designs that facilitate robust DE reformation from irregular SE streams that can result from manipulations on released DE cores. We identified operational parameters and flow regimes capable of reforming high-quality DEs from the manipulated droplets to obtain useful DE outputs. By combining core-release and DE reforming strategies, it is now possible to effectively switch emulsion formats on demand. This allows for the benefits of DEs in terms of coalescence properties and screening with flow-cytometry to integrate with SE strengths in on-chip manipulations. These strategies enabled a novel multistep DE aptamer screen to be performed, demonstrating the potential for new DE-based assays leveraging multistep reactions. This work advances the functionality of double emulsion-based assays to reach or surpass the level of single emulsion-based assays by enabling active manipulations in a new drop format with unique emulsion properties.

## Acknowledgements

This work was carried out in part in the Materials Research Laboratory Central Research Facilities, University of Illinois. We would like to thank the Roy J. Carver Biotechnology Center for their support. We also thank the University of Illinois at Urbana-Champaign for funding through the UIUC Startup Grant. We further recognize financial support from the Gordon and Betty Moore Foundation through Grant GBMF9195 to the Carl R. Woese Institute of Genomic Biology.

## Author Contributions

T.W.C. and H.-S.H designed experiments. T.W.C. and W.J. designed and fabricated microfluidic devices. T.W.C. performed droplet experiments. T.W.C., W.J., and H.-S.H prepared the manuscript.

---

**Scheme 1:**
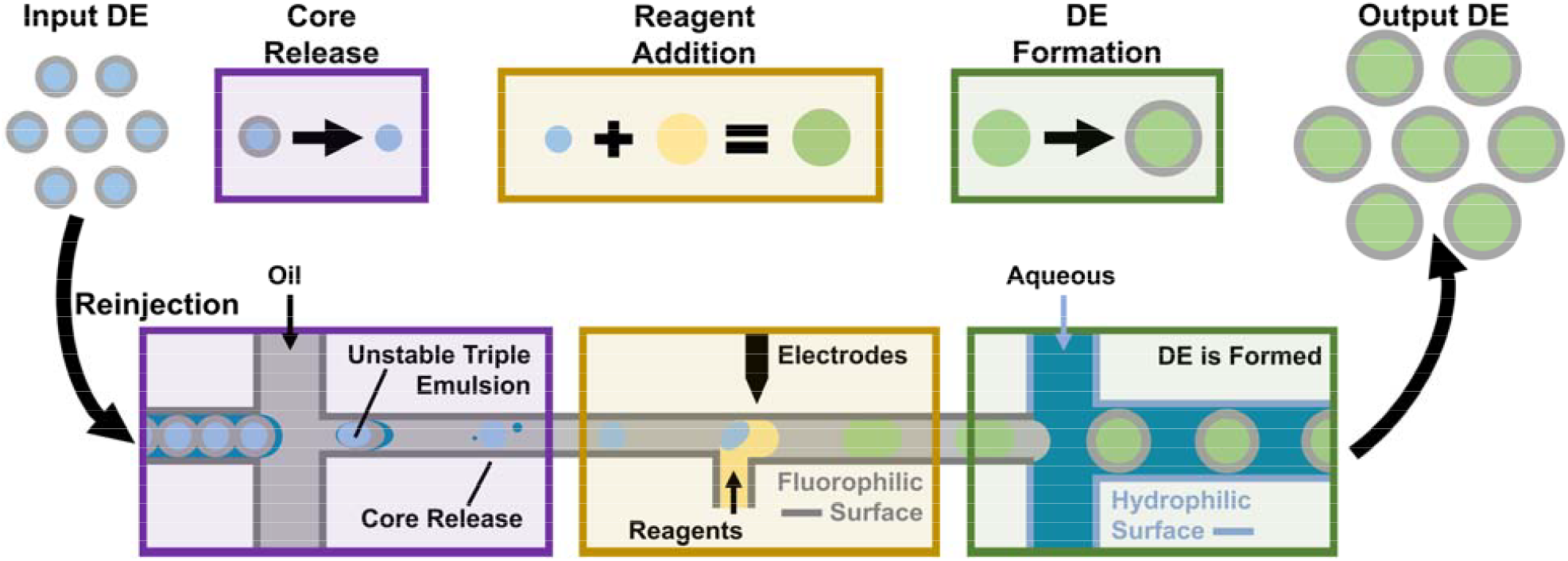
A) Schematic representation of reagent addition in DEs. A monodisperse DE input is reinjected and respaced then converted to a SE on-chip through core release. The SE drops then undergo reagent addition by picoinjection. The resulting droplets are dispersed in water to reform a DE output.

**Scheme 2:**
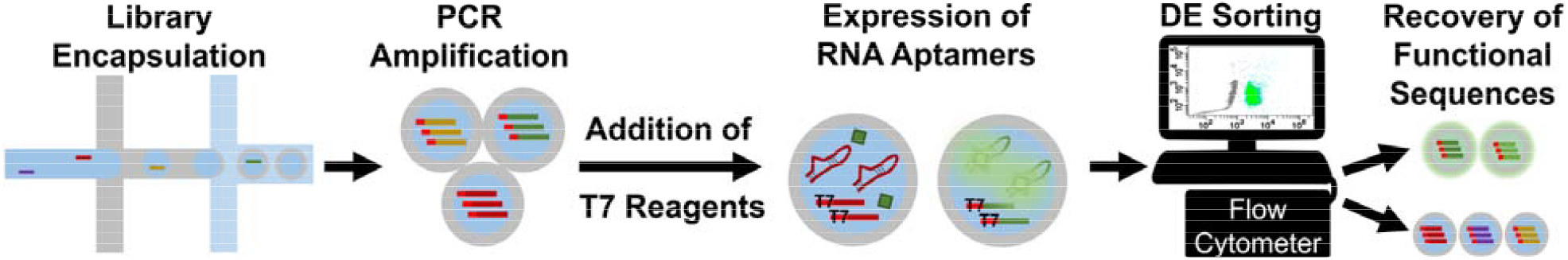
Workflow for a double emulsion screen of a ssDNA library encoding RNA light-up aptamers. Digital amplification occurs in DEs, followed by reagent addition and digital RNA expression in the DE. DE-Flow Cytometry recovers fluorescent light-up aptamer sequences.

## Notes

### Competing Interest Statement

The authors have declared no competing interest.

## References

1. Teh, S. Y., Lin, R., Hung, L. H. & Lee, A. P. Droplet microfluidics. Lab on a Chip vol. 8 198–220 Preprint at 10.1039/b715524g (2008).

2. Niculescu, A. G., Chircov, C., Bîrcă, A. C. & Grumezescu, A. M. Fabrication and applications of microfluidic devices: A review. International Journal of Molecular Sciences vol. 22 1–26 Preprint at 10.3390/ijms22042011 (2021).

3. Abatemarco, J. et al. RNA-aptamers-in-droplets (RAPID) high-throughput screening for secretory phenotypes. Nat Commun 8, 1–9 (2017).

4. Hatori, M. N., Modavi, C., Xu, P., Weisgerber, D. & Abate, A. R. Dual-layered hydrogels allow complete genome recovery with nucleic acid cytometry. Biotechnol J 17, 2100483 (2022).

5. Zhu, Z. et al. Single-molecule emulsion PCR in microfluidic droplets. Analytical and Bioanalytical Chemistry vol. 403 2127–2143 Preprint at 10.1007/s00216-012-5914-x (2012).

6. Stucki, A., Vallapurackal, J., Ward, T. R. & Dittrich, P. S. Droplet Microfluidics and Directed Evolution of Enzymes: An Intertwined Journey. Angewandte Chemie - International Edition 60, 24368–24387 (2021).

7. Fang, X. et al. Rapid screening of aptamers for fluorescent targets by integrated digital PCR and flow cytometry. Talanta 242, 123302 (2022).

8. Zilionis, R. et al. Single-cell barcoding and sequencing using droplet microfluidics. Nat Protoc 12, 44–73 (2017).

9. Payne, E. M., Holland-Moritz, D. A., Sun, S. & Kennedy, R. T. High-throughput screening by droplet microfluidics: perspective into key challenges and future prospects. Lab Chip 20, 2247–2262 (2020).

10. Tabuchi, T. & Yokobayashi, Y. High-throughput screening of cell-free riboswitches by fluorescence-activated droplet sorting. Nucleic Acids Res 50, 3535–3550 (2022).

11. Moragues, T. et al. Droplet-based microfluidics. Nature Reviews Methods Primers 3, 32 (2023).

12. Utada, A. S. et al. Monodisperse Double Emulsions Generated from a Microcapillary Device. Science (1979) 308, 537–541 (2005).

13. Anna, S. L., Bontoux, N. & Stone, H. A. Formation of dispersions using “flow focusing” in microchannels. Appl Phys Lett 82, 364–366 (2003).

14. Shah, R. K. et al. Designer emulsions using microfluidics. Materials Today vol. 11 18–27 Preprint at 10.1016/S1369-7021(08)70053-1 (2008).

15. Kim, S.-H., Kim, J. W.Cho, J.-C. & Weitz, D. A. Double-emulsion drops with ultra-thin shells for capsule templates. Lab Chip 11, 3162–3166 (2011).

16. Lee, T. Y., Choi, T. M., Shim, T. S., Frijns, R. A. M. & Kim, S. H. Microfluidic production of multiple emulsions and functional microcapsules. Lab on a Chip vol. 16 3415–3440 Preprint at 10.1039/c6lc00809g (2016).

17. Fattahi, P. et al. Core–shell hydrogel microcapsules enable formation of human pluripotent stem cell spheroids and their cultivation in a stirred bioreactor. Sci Rep 11, 7177 (2021).

18. Calhoun, S. G. K. et al. Systematic characterization of effect of flow rates and buffer compositions on double emulsion droplet volumes and stability. Lab Chip 22, 2315–2330 (2022).

19. Brower, K. K. et al. Double emulsion flow cytometry with high-throughput single droplet isolation and nucleic acid recovery. Lab Chip 20, 2062–2074 (2020).

20. Cowell, T. W., Dobria, A. & Han, H.-S. Simplified, Shear Induced Generation of Double Emulsions for Robust Compartmentalization during Single Genome Analysis. ACS Appl Mater Interfaces 14, 20528–20537 (2022).

21. Brower, K. K. et al. Double Emulsion Picoreactors for High-Throughput Single-Cell Encapsulation and Phenotyping via FACS. Anal Chem 92, 13262–13270 (2020).

22. McCully, A. L., Loop Yao, M., Brower, K. K., Fordyce, P. M. & Spormann, A. M. Double emulsions as a high-throughput enrichment and isolation platform for slower-growing microbes. ISME Communications 3, 1–9 (2023).

23. Ma, S., Huck, W. T. S. & Balabani, S. Deformation of double emulsions under conditions of flow cytometry hydrodynamic focusing. Lab Chip 15, 4291–4301 (2015).

24. Lim, S. W. & Abate, A. R. Ultrahigh-throughput sorting of microfluidic drops with flow cytometry. Lab Chip 13, 4563–4572 (2013).

25. Nuti, N. et al. A Multiplexed Cell-Free Assay to Screen for Antimicrobial Peptides in Double Emulsion Droplets. Angewandte Chemie - International Edition 61, e202114632 (2022).

26. Cowell, T. & Han, H.-S. Double Emulsion Flow Cytometry for Rapid Single Genome Detection. Methods in molecular biology 2689, 155–167 (2023).

27. Baret, J.-C. et al. Fluorescence-activated droplet sorting (FADS): efficient microfluidic cell sorting based on enzymatic activity. Lab Chip 9, 1850 (2009).

28. Mazutis, L. et al. Single-cell analysis and sorting using droplet-based microfluidics. Nat Protoc 8, 870–891 (2013).

29. Cole, R. H. et al. Printed droplet microfluidics for on demand dispensing of picoliter droplets and cells. Proceedings of the National Academy of Sciences 114, 8728–8733 (2017).

30. Adan, A., Alizada, G., Kiraz, Y., Baran, Y. & Nalbant, A. Flow cytometry: basic principles and applications. Crit Rev Biotechnol 37, 163–176 (2017).

31. Robinson, J. P., Ostafe, R., Iyengar, S. N., Rajwa, B. & Fischer, R. Flow Cytometry: The Next Revolution. Cells vol. 12 1875 Preprint at 10.3390/cells12141875 (2023).

32. Sukovich, D. J., Lance, S. T. & Abate, A. R. Sequence specific sorting of DNA molecules with FACS using 3dPCR. Sci Rep 7, 39385 (2017).

33. McCully, A. L., Loop Yao, M., Brower, K. K., Fordyce, P. M. & Spormann, A. M. Double emulsions as a high-throughput enrichment and isolation platform for slower-growing microbes. ISME Communications 3, 1–9 (2023).

34. Mazutis, L. et al. Single-cell analysis and sorting using droplet-based microfluidics. Nat Protoc 8, 870–891 (2013).

35. Klein, A. M. et al. Droplet barcoding for single-cell transcriptomics applied to embryonic stem cells. Cell 161, 1187–1201 (2015).

36. Habib, N. et al. Massively parallel single-nucleus RNA-seq with DroNc-seq. Nat Methods 14, 955–958 (2017).

37. De Rop, F. V. et al. HyDrop enables droplet based single-cell ATAC-seq and single-cell RNA-seq using dissolvable hydrogel beads. Elife 11, (2022).

38. Hindson, B. J. et al. High-Throughput Droplet Digital PCR System for Absolute Quantitation of DNA Copy Number. Anal Chem 83, 8604–8610 (2011).

39. Abate, A. R., Hung, T., Mary, P., Agresti, J. J. & Weitz, D. A. High-throughput injection with microfluidics using picoinjectors. Proceedings of the National Academy of Sciences 107, 19163–19166 (2010).

40. Sciambi, A. & Abate, A. R. Generating electric fields in PDMS microfluidic devices with salt water electrodes. Lab Chip 14, 2605–2609 (2014).

41. Priest, C., Herminghaus, S. & Seemann, R. Controlled electrocoalescence in microfluidics: Targeting a single lamella. Appl Phys Lett 89, 134101 (2006).

42. Hou, L. et al. Continuously Electrotriggered Core Coalescence of Double-Emulsion Drops for Microreactions. ACS Appl Mater Interfaces 9, 12282–12289 (2017).

43. Jia, Y. et al. Sequential Coalescence Enabled Two-Step Microreactions in Triple-Core Double-Emulsion Droplets Triggered by an Electric Field. Small 13, (2017).

44. Qu, F. et al. Thermo-Induced Coalescence of Dual Cores in Double Emulsions for Single-Cell RT-PCR. Anal Chem 94, 11670–11678 (2022).

45. Chen, X. et al. NIR light-triggered core-coalescence of double-emulsion drops for micro-reactions. Chemical Engineering Journal 454, 140050 (2023).

46. Stucki, A., Jusková, P., Nuti, N., Schmitt, S. & Dittrich, P. S. Synchronized Reagent Delivery in Double Emulsions for Triggering Chemical Reactions and Gene Expression. Small Methods 5, 2100331 (2021).

47. Lee, S. S., Abbaspourrad, A. & Kim, S. H. Nonspherical double emulsions with multiple distinct cores enveloped by ultrathin shells. ACS Appl Mater Interfaces 6, 1294–1300 (2014).

48. Elvira, K. S., Gielen, F., Tsai, S. S. H. & Nightingale, A. M. Materials and methods for droplet microfluidic device fabrication. Lab Chip 22, 859–875 (2022).

49. Sciambi, A. & Abate, A. R. Adding reagent to droplets with controlled rupture of encapsulated double emulsions. Biomicrofluidics 7, 044112 (2013).

50. Abate, A. R., Hung, T., Mary, P., Agresti, J. J. & Weitz, D. A. High-throughput injection with microfluidics using picoinjectors. Proceedings of the National Academy of Sciences 107, 19163–19166 (2010).

51. Filonov, G. S., Moon, J. D., Svensen, N. & Jaffrey, S. R. Broccoli: Rapid selection of an RNA mimic of green fluorescent protein by fluorescence-based selection and directed evolution. J Am Chem Soc 136, 16299–16308 (2014).

52. Sun, Z. et al. Live-Cell Imaging of Guanosine Tetra- and Pentaphosphate (p)ppGpp with RNA-based Fluorescent Sensors*. Angew Chem Int Ed Engl 60, 24070–24074 (2021).

53. Peng, Y. et al. Live-Cell Imaging of Endogenous RNA with a Genetically Encoded Fluorogenic Allosteric Aptamer. Anal Chem 95, 13762–13768 (2023).

54. Han, K. Y., Leslie, B. J., Fei, J., Zhang, J. & Ha, T. Understanding the photophysics of the spinach-DFHBI RNA aptamer-fluorogen complex to improve live-cell RNA imaging. J Am Chem Soc 135, 19033–8 (2013).

55. Wang, J. et al. Particle display: a quantitative screening method for generating high-affinity aptamers. Angew Chem Int Ed Engl 53, 4796–801 (2014).

56. Ryckelynck, M. et al. Using droplet-based microfluidics to improve the catalytic properties of RNA under multiple-turnover conditions. RNA 21, 458–469 (2015).

57. Filonov, G. S., Song, W. & Jaffrey, S. R. Spectral Tuning by a Single Nucleotide Controls the Fluorescence Properties of a Fluorogenic Aptamer. Biochemistry 58, 1560–1564 (2019).

58. Chen, X., Brukson, A. & Ren, C. L. A simple droplet merger design for controlled reaction volumes. Microfluid Nanofluidics 21, 34 (2017).

59. Link, D. R., Anna, S. L., Weitz, D. A. & Stone, H. A. Geometrically Mediated Breakup of Drops in Microfluidic Devices. Phys Rev Lett 92, 054503 (2004).

60. Nie, J. & Kennedy, R. T. Sampling from nanoliter plugs via asymmetrical splitting of segmented flow. Anal Chem 82, 7852–7856 (2010).

61. Bauer, W. A. C., Fischlechner, M., Abell, C. & Huck, W. T. S. Hydrophilic PDMS microchannels for high-throughput formation of oil-in-water microdroplets and water-in-oil-in-water double emulsions. Lab Chip 10, 1814–1819 (2010).

